# Simulation of model overfit in variance explained with genetic data

**DOI:** 10.1101/598904

**Authors:** Jaime Derringer

## Abstract

Two recent papers, and an author response to prior commentary, addressing the genetic architecture of human temperament and character claimed that “The identified SNPs explained nearly all the heritability expected”. The authors’ method for estimating heritability may be summarized as: Step 1: Pre-select SNPs on the basis of GWAS p<0.01 in the target sample. Step 2: Enter target sample genotypes (the pre-selected SNPs from Step 1) and phenotypes into an unsupervised machine learning algorithm (Phenotype-Genotype Many-to-Many Relations Analysis, PGMRA) for further reduction of the set of SNPs. Step 3: Test the sum score of the SNPs identified from Step 2, weighted by the GWAS regression weights estimated in Step 1, within the same target sample. The authors interpreted the linear regression model R^2^ obtained from Step 3 as a measure of successfully identified heritability. Regardless of the method applied to select SNPs in Step 2, the combination of Steps 1 and 3, as described, causes inflation of the estimated effect size. The extent of this inflation is demonstrated here, where random SNP selection and polygenic scoring from simulated random data recovered effect sizes similar to those reported in the original empirical papers.

## Background

An author response [1] to my previous commentary [2] claimed that the application of PGMRA in parallel papers on the genetics of character [4] and temperament [5] was not subject to concerns about overfitting. Unlike polygenic scoring, the goal was not to identify a generalizable combination of genetic variants to apply to outside samples. Rather, the stated goal was to estimate the heritability in the phenotype that could be attributed to the identified SNPs. The method for estimating what was interpreted as heritability (as illustrated in the author response, Figure 1 [1]) may be summarized as:

**Figure 1.**
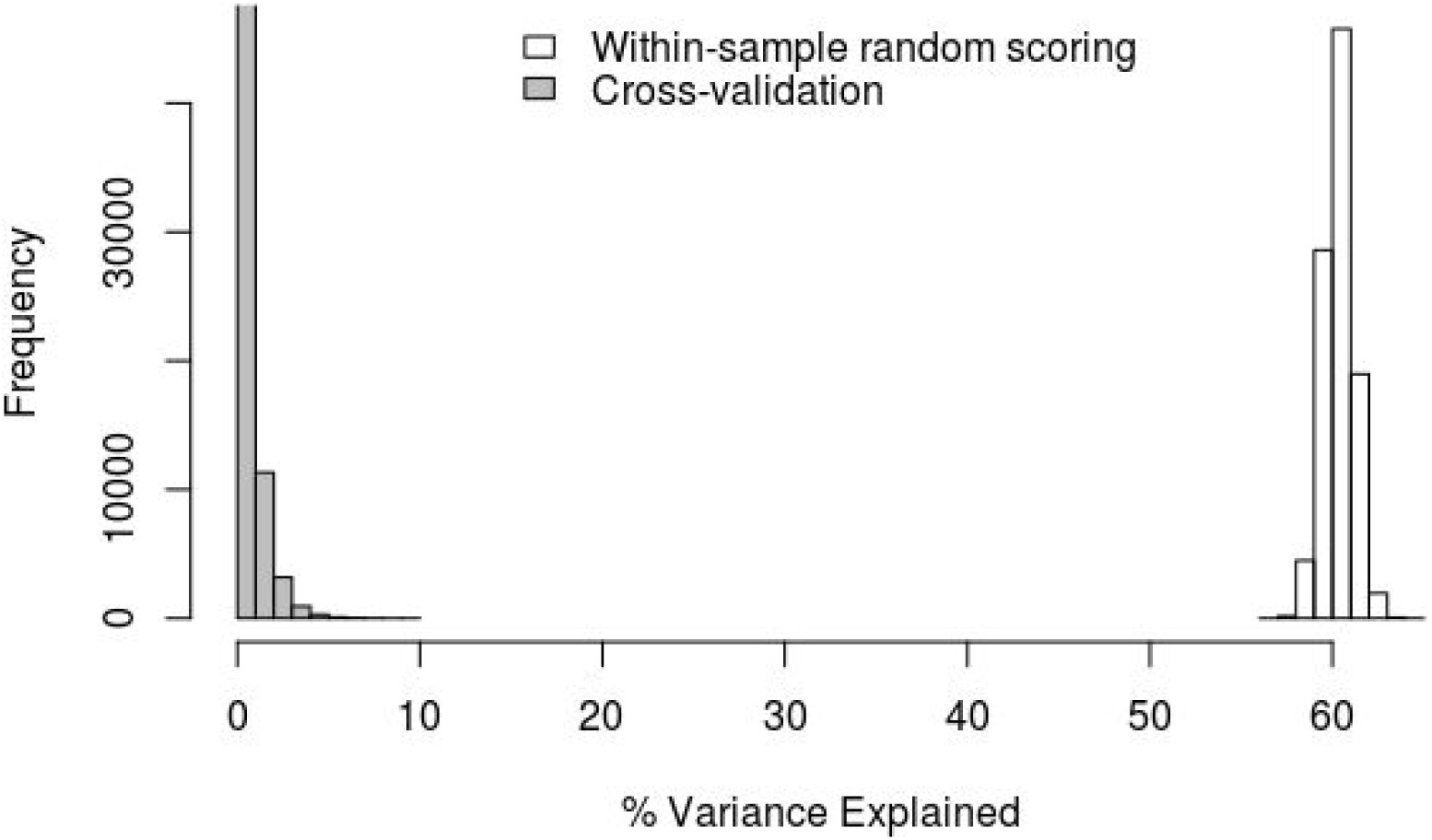
R^2^ from 100 000 simulations of random sets of 359 SNPs scored and applied within the discovery sample (N = 2 000, white bars) versus cross-validated in an independent replication sample (N = 200, grey bars). Full script available at: https://osf.io/t5jbs/.

Step 1: Pre-select SNPs on the basis of GWAS p<0.01 in the target sample.

Step 2: Enter target sample genotypes (the pre-selected SNPs from Step 1) and phenotypes into PGMRA [3] for further reduction of the set of SNPs.

Step 3: Test the sum score of the SNPs identified from Step 2, weighted by the GWAS regression weights estimated in Step 1, within the same target sample.

The authors interpreted the linear regression model R^2^ obtained from Step 3 as a measure of successfully identified heritability. Regardless of the method applied to select SNPs in Step 2 - whether it is global or local, data-driven [4,5] or based on theory [6,7] - the combination of Steps 1 and 3, as described, causes inflation of the estimated effect size.

## Method

To demonstrate the extent of effect size inflation resulting from the above-described method, I present a simulation in which random genotypes and phenotypes were generated and Steps 1 and 3 were applied as described by the authors [1,4,5]. Here, however, Step 2 was instead executed as random selection of SNPs for inclusion in the score. The entire process, plus cross-validation for comparison, was performed 1000 times per each of 100 simulated data sets, each comprised of 700 000 SNPs, with discovery and replication sample sizes of 2 000 and 200, respectively. Given that the genotypes, phenotypes, and selected SNPs were entirely random, a correct estimate of the effect should be essentially zero, and any deviation from zero in the estimate of variance explained may be taken as an indicator of model bias, not true heritability.

The extent of inflation in the score is dependent on the number of included predictors. For the current simulation, the number of SNPs included in each random score was set equal to the number of SNPs remaining after pruning for linkage disequilibrium (LD, using a restrictive threshold of r^2^ > 10%) the SNPs identified by PGMRA in the largest sample in the character paper (Finns, N = 2 149 [4]). LD structure was estimated in the 1000 Genomes phase 3 Finnish (FIN) reference population [8]. This process yielded an estimate of 359 independent SNPs as an appropriate comparison to the original paper results [4]. The full script for this simulation is available at: https://osf.io/t5jbs/. Simulation and analysis used R [9] and Plink [10].

## Result

The result of these simulations are presented in Figure 1. Within sample scoring, where the identification of SNPs and their effects was estimated and tested within a single sample as described [1,4,5] yielded an average estimate of 60% variance explained (SD = 0.008). This is not less than the variance explained reported by the authors in their non-random datasets (57% and 48% for the Finnish samples applied to character and temperament, respectively [4,5]). When the same randomly selected scores were applied in a cross-validated manner (that is, identified within one sample, tested in an another), the effect size essentially (and correctly, given the random nature of the data) dropped to zero (M = 0.5%, max = 9%). Although the authors noted that such cross-validated effect sizes may be reduced in the presence of heterogeneity across samples [1], the current simulation generated the sets of genotypes and phenotypes for the discovery and replication samples simultaneously, so that systematic or structural between-sample heterogeneity would not be a likely explanation for the reduction in effect size demonstrated here.

## Conclusion

The observed inflation of the effect size is caused by the combination of Steps 1 and 3, where predictors were selected and tested within the same sample. In no way is this the result of unique properties of any additional selection procedures that may take place in Step 2 (whether PGMRA [3] or otherwise), nor is model overfitting in this manner a problem that is unique to issues of genetic data or machine learning. Although technical capacities and computational models may increase in complexity, such advances do not necessarily overcome the basic requirements of model estimation and evaluation in general. The current simulation does not explore the potential utility of PGMRA for identifying promising sets of associated SNPs. However, the extent of inflation in the described estimate of “heritability” (the within-sample variance explained; as reported in the abstracts of both original papers [4,5]) is so substantial as to be uninterpretable.

